# Cellular Senescence Affects ECM Regulation in COPD Lung Tissue

**DOI:** 10.1101/2023.08.02.551614

**Authors:** R.R. Woldhuis, N.J. Bekker, J.L.L. van Nijnatten, M. Banchero, W. Kooistra, J.C. Wolters, P.L. Horvatovich, V. Guryev, M. van den Berge, W. Timens, C.A. Brandsma

**Affiliations:** Department of Pathology and Medical Biology, University of Groningen, University Medical Centre Groningen; Groningen Research Institute for Asthma and COPD (GRIAC), University of Groningen, University Medical Centre Groningen; Department of Pediatrics, University of Groningen, University Medical Centre Groningen; Department of Analytical Biochemistry, University of Groningen, University Medical Centre Groningen; European Research Institute for the Biology of Aging (ERIBA), University of Groningen, University Medical Centre Groningen; Department of Pulmonary Diseases, University of Groningen, University Medical Centre Groningen; University of Technology Sydney, Respiratory Bioinformatics and Molecular Biology, Sydney, New South Wales, Australia

**Keywords:** COPD, Cellular senescence, extracellular matrix, ECM dysregulation, proteases

## Abstract

**Introduction:** Higher levels of cellular senescence have been demonstrated in COPD patients, including severe early-onset (SEO)-COPD. Recently, we demonstrated that senescence induces extracellular matrix (ECM) dysregulation in lung fibroblasts. However, this *in vitro* observation has not been demonstrated *in vivo* yet. Therefore, we investigated whether cellular senescence can contribute to COPD-associated ECM changes in parenchymal lung tissue.

**Methods:** Transcriptomic and proteomic analyses were performed on parenchymal lung tissue from 60 COPD patients (including 18 SEO-COPD patients) and 32 controls. Differential expression of ECM-related genes and proteins was compared between (SEO-)COPD and controls, followed by correlations with six senescence markers and four senescence signature scores. Significant ECM-senescence correlations were verified using histology and primary lung fibroblasts.

**Results:** We identified 12 COPD- and 57 SEO-COPD-associated ECM genes and 4 COPD- and 9 SEO-COPD-associated ECM proteins of which the majority, 45 genes and 5 proteins, correlated with senescence. The correlations for *COL6A1, COL6A2* and *FBLN5* were confirmed in situ and correlations for 21 ECM genes were confirmed in primary lung fibroblasts at baseline. Four genes were successfully functionally validated in our senescence-induced lung fibroblast model, including increased protein levels of ADAMST1 and a non-functional FBLN5 protein.

**Conclusions:** We confirm a strong link between (SEO-)COPD-associated ECM changes and senescence *in vivo* in peripheral lung tissue from COPD patients. The strongest and most consistent senescence-associated ECM components include proteases, elastogenesis genes, and collagens 6. These results indicate a contributing role for senescence in disturbed ECM and elastic fiber organization, and protease-antiprotease imbalance in COPD.

## INTRODUCTION

Cellular senescence is a major ageing hallmark demonstrated in COPD patients, both in lung tissue, and structural and immune cells ^(1, 2)^. Cellular senescence is defined by irreversible cell cycle arrest. Senescent cells have an impaired cellular function and secrete a panel of pro-inflammatory chemokines, cytokines, growth factors, and proteases, called ‘senescence-associated secretory phenotype’ (SASP). This SASP causes chronic inflammation in the surrounding tissue and contributes to tissue dysfunction and remodeling ^(3, 4)^.

Another hallmark of lung ageing and COPD is extracellular matrix (ECM) dysregulation ^(5)^. The ECM is essential for the function and structure of the lungs and is involved in healthy lung tissue repair. Lung fibroblasts are major producers of ECM and regulate ECM homeostasis. In the parenchyma of COPD patients, reduction and breakdown of elastic fibers and other ECM proteins are observed ^(6, 7)^. In addition, reduced levels of proteoglycans involved in the organization of collagen crosslinking are found, including decorin and biglycan ^(8)^. Furthermore, the protease-antiprotease balance is disturbed in COPD, which causes breakdown of the ECM mainly in the parenchyma eventually resulting in emphysema ^(9)^. Age-related ECM changes can affect normal ECM homeostasis and healthy tissue function ^(10)^, including loss of elasticity ^(7, 11)^, similar to what is observed in COPD. This indicates a role for ageing in ECM dysregulation in COPD patients.

Recently, we demonstrated higher levels of cellular senescence in parenchymal lung fibroblasts derived from COPD patients compared to matched non-COPD controls, which was most pronounced in fibroblasts from severe early-onset (SEO-)COPD patients ^(12)^. These SEO-COPD patients are of particular interest as they have (very) severe airflow obstruction (FEV1 < 40% predicted) at relatively early age (before 55 years of age) and therefore seem to be very susceptible to COPD development. In this previous study, we also found a direct link between higher levels of senescence and ECM dysregulation, with lower levels of regulators of ECM organization and structural ECM proteins, including decorin and biglycan^(12)^.

Since we previously demonstrated that cellular senescence leads to ECM dysregulation *in vitro*, the next step is to investigate whether this association is also present *in vivo*. Therefore, we first identified the COPD-associated ECM changes in lung tissue using comprehensive transcriptomic and proteomic analyses and next assessed which of these ECM changes are associated with senescence. Significant ECM-senescence correlations in lung tissue were verified in situ using histology and on cellular level using primary lung fibroblasts.

## MATERIALS & METHODS

*A full description of the methods can be found in the online supplement*.

### Lung tissue collection for transcriptomic and proteomic analyses

Peripheral lung tissue from current-smoking and ex-smoking non-COPD controls (FEV_1_/FVC>70% of predicted) and COPD (FEV_1_/FVC<70% of predicted) patients (including SEO-COPD; FEV_1_<40% of predicted and age≤55) was used from the PRoteogenomics Early-onSeT cOpd (PRESTO) lung cohort (described in supplement). Transcriptomic and proteomic data from 60 COPD patients and 32 controls was used (see table 1 for characteristics).

**Table 1:** Patient characteristics of lung tissue subjects for transcriptomic and proteomic analyses.

| Variable | Non-COPD (ex-)smoker control | All COPD patients | Subgroup of SEO-COPD patients |
| --- | --- | --- | --- |
| Number | 32 | 60 | 18 |
| Age, median (range) | 62 (45-76) | 59 (39-79) | 52 (39-55) # |
| Male/female, N | 15/17 | 31/29 | 5/13 |
| Smoking status (ES/CS) | 13/19 | 55/5 * | 18/0 # |
| Pack-years, median (range) | 35 (10-75) | 32 (6-130) | 28 (6-54) # |
| non-COPD, N | 32 | - | - |
| COPD, N | - | 60 | 18 |
| GOLD 1 | - | - | - |
| GOLD 2 | - | 14 | - |
| GOLD 3 | - | 16 | 2 |
| GOLD 4 | - | 30 | 16 |
| FEV<sub>1</sub> %pred, median (range) | 85.0 (59.8-125.0) | 30.5 (10.6-77.0) * | 17.3 (12.0-37.0) # |
| FEV<sub>1</sub>/FVC, median (range) | 76.9 (71.0-86.6) | 41.6 (18.8-67.9) * | 29.2 (20.5-50.0) # |

ES = ex-smoker, CS = current smoker, FEV<sub>1</sub> = forced expiratory volume in the first second, FVC = Forced vital capacity, %pred = percentage of predicted. Mann-Whitney U tests or Chi-square tests were used to test differences in characteristics for COPD patients and SEO-COPD patients compared to non-COPD controls and indicated (\* for COPD and # for SEO-COPD) when significant ( $p < 0.05$ ). For transcriptomics analyses, 4 COPD samples did not pass QC and were therefore excluded for analyses, which did not affect the statistics.

### Lung tissue transcriptomics

RNA sequencing was performed on RNA derived from frozen lung tissue sections. A list of 471 ECM-related genes based on three Gene Ontology databases (GO:0085029, GO:0062023, GO:0005201) was used for differential expression analyses. ECM gene expression was compared between control current smokers (CS) and control ex-smokers (ES) first to exclude genes affected by smoking status. Next, ECM gene expression was compared between (SEO-)COPD and control using generalized linear models corrected for age and sex. The (SEO-)COPD-associated ECM genes were correlated with six major senescence genes (*CDKN2A, CDKN1A, TP53, CDKN2B, CDKN1B*, and *LMNB1*) and the calculated GSVA scores of four senescence signatures (SenMayo ^(13)^, Casella ^(14)^, Hernandez-Segura ^(15)^, and Fridman ^(16)^) in all available lung tissue samples. Benjamini-Hochberg False discovery rate (FDR) below 0.05 was considered statistically significant. Meta-analyses were performed to identify ECM genes with consistent correlations with multiple senescence genes and signature scores. P-value below 0.05 was considered statistically significant.

### Lung tissue proteomics

Serial sections from the same lung tissue frozen section samples as used for transcriptomics, were used for discovery-based mass spectrometry proteomics. Protein levels of the selected ECM-related proteins were compared between control current smokers (CS) and control ex-smokers (ES) first to exclude proteins affected by smoking status. Next, ECM protein levels were compared between (SEO-)COPD and control using generalized linear model corrected for age and sex. The (SEO-)COPD-associated ECM proteins were correlated with the six major senescence genes and the GSVA scores of the four senescence signatures in all available lung tissue samples. Benjamini-Hochberg False discovery rate (FDR) below 0.05 was considered statistically significant.

### Immunohistochemical verification on lung tissue

Data from the HOLLAND study regarding COL6A1, COL6A2, ELN, FBLN5, and LUM staining in lung tissue, described by Joglekar et al. ^(17)^, and p21 staining, described by Chen et al. ^(18)^, was used to verify ECM-senescence correlations in situ using lung tissue sections from ex-smoking COPD patients (n=27) and ex-smoking non-COPD controls (n=18). Area of positive p21 staining was correlated with area of positive ECM staining and intensity of ECM staining in the parenchymal and airway wall regions. For verification, P-value below 0.05 was considered statistically significant.

### Primary lung fibroblast culture

Primary lung fibroblasts from SEO-COPD (n=10) and non-COPD patients (n=11) (see Table E1 for patient characteristics) were cultured at baseline conditions and cellular senescence was induced in non-COPD-derived fibroblasts using our previously described Paraquat-induced senescence model ^(12)^. The (SEO-)COPD- and senescence-associated ECM genes were confirmed in fibroblasts using RNA sequencing, and the effect of senescence induction was assessed using RNA sequencing, ELISA, and Western Blot for Fibulin-5. For confirmation, P-value below 0.05 was considered statistically significant.

## RESULTS

### Patient characteristics

The clinical characteristics of the patients included from the PRESTO cohort are shown in Table 1. Smoking status was significantly different between (SEO-)COPD and control. Therefore, we first excluded differentially expressed (p<0.05) genes and proteins between current and ex-smoking controls. In addition, age and pack-years of smoking were significantly different between SEO-COPD and control. Therefore, we corrected for age to account for these (highly correlated) confounders.

### (SEO-)COPD-associated ECM-related genes and proteins

Of the 471 ECM-related proteins selected from the Gene Ontology databases, 354 genes and 190 proteins were detected in lung tissue. On transcript level, 12 ECM genes were differentially expressed (FDR<0.05) between COPD-derived and control-derived lung tissue (Figure 1A). On protein level, 4 ECM proteins were different between COPD and control (Figure 1B).

**Figure 1:**
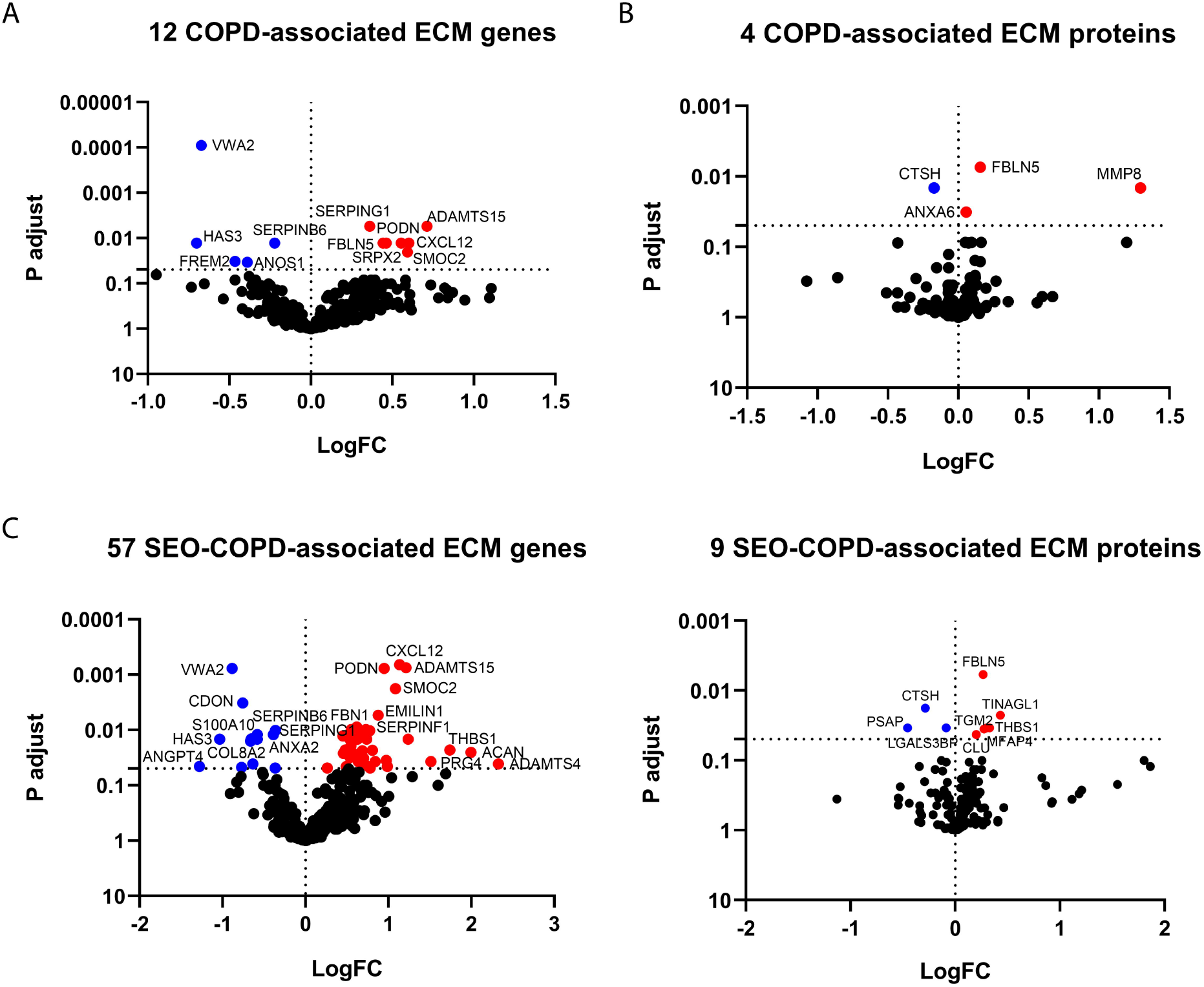
(SEO-)COPD-associated ECM-related genes and proteins in lung tissue. ECM-related gene expression and protein levels were compared between COPD and controls (A+B) and SEO-COPD and controls (C+D) using transcriptomic (A+C) and proteomic (B+D) analyses. Volcano plots depict the LogFC (X-axis) and FDR (P adjust, Y-axis) of all detected ECM-related genes and proteins. Significant (P adjust < 0.05) differentially expressed ECM-related genes and proteins are depicted in color, with higher expression in COPD in red and lower expression in COPD in blue. Horizontal dotted lines represent significance cut-off (FDR = 0.05), and vertical dotted lines represent LogFC of 0.

In the SEO-COPD subgroup, 57 ECM genes were differentially expressed compared to control (Figure 1C). All 12 COPD-associated ECM genes were also significantly different in the same direction in SEO-COPD. On protein level, 9 ECM proteins were different between SEO-COPD and control (Figure 1D).

### (SEO-)COPD-associated ECM-related genes and proteins are correlated with senescence

To investigate the association between ECM and senescence, all significant (SEO-)COPD-associated ECM genes and proteins were correlated with six well-known senescence genes (*CDKN1A/p21, CDKN1B/p27, CDKN2A/p16, CDKN2B/p15, TP53*, (up-regulated with senescence) and *LMNB1* (down-regulated with senescence)). Of the 57 ECM genes, 37 were correlated with one or more of the 6 senescence genes (Figure 2A), including 26 genes correlating with CDKN1A/p21, of which all positively correlated genes were higher expressed in SEO-COPD and all negatively correlated genes were lower expressed in SEO-COPD. In addition, 17 ECM genes were correlated with *LMNB1*, of which 11 genes were positively correlated. Of the 11 (SEO-)COPD-associated ECM proteins, 5 were positively correlated with *CDKN1A*/p21, which were all higher expressed in (SEO-)COPD (Figure 2B).

**Figure 2:**
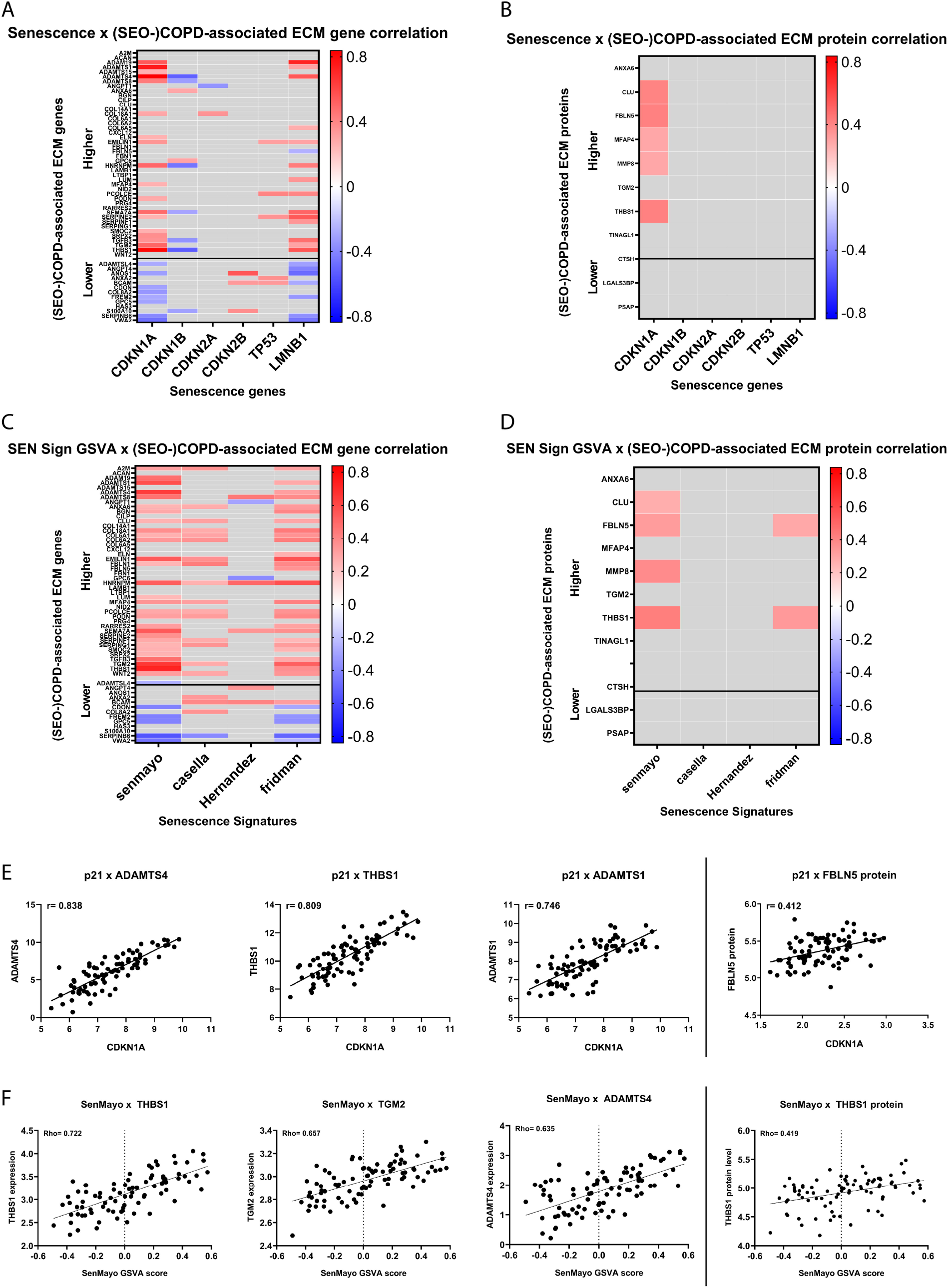
Correlations of senescence genes and GSVA scores of senescence signatures with (SEO-)COPD-associated ECM-related genes and proteins. All (SEO-)COPD associated ECM-related genes (A+C) and proteins (B+D) were correlated with the six major senescence genes using Pearson’s correlations (A+B) and with the GSVA scores of the four senescence signatures using Spearman’s correlations (C+D). The heatmaps depict the significant correlations (FDR < 0.05) in color and the non-significant correlations in grey. The colors represent the strength (r or rho coefficient) of the correlations, with positive correlations in red and negative correlations in blue. The ECM-related genes and proteins higher expressed in COPD are depicted on top and the ones lower expressed in COPD are depicted on the bottom of the y-axis. The senescence genes and GSVA scores of senescence signatures are depicted on the X-axis. The strongest (SEO-)COPD-associated ECM and senescence correlations are shown the scatter plots with senescence gene correlations on top (E) and senescence signature correlations on the bottom (F). The Pearson’s or Spearman’s correlation coefficients (r or rho) are depicted in the top left corner of the graphs.

Next, the (SEO-)COPD-associated ECM genes were correlated with GSVA scores of four commonly used senescence signatures. 43 out of 57 ECM genes were correlated with one or more senescence scores (Figure 2C), with 31 genes correlating with scores of at least 2 signatures. Most ECM genes correlated with the SenMayo and Fridman scores, which were both higher expressed in SEO-COPD compared to control lung tissue (Figure E1). Importantly, 24 of 26 *CDKN1A*/p21-correlated ECM genes were also correlated with a senescence score, strengthening the ECM-senescence correlations in lung tissue. On protein level, 4 out of 11 ECM proteins were correlated with one or more senescence scores (Figure 2D).

The three strongest *CDKN1A*/p21 correlated genes are *ADAMTS4, THBS1*, and *ADAMTS1* (Figure 2E), of which *THBS1* and *ADAMTS4* were also among the three strongest ECM genes correlated with a senescence score (Figure 2F). FBLN5 and THBS1 protein levels were both correlated with *CDKN1A*/p21 and SenMayo score (Figure 2E+F).

### Meta-analyses of correlations reveal consistent senescence-associated ECM genes

Meta-analyses were performed on the five up-regulated senescence genes and the four senescence scores to assess the consistency of the ECM-senescence correlations across the different senescence markers. Seven ECM genes were consistently associated with the up-regulated senescence genes (Figure 3A). 33 ECM genes were consistently associated with the four senescence scores (Figure 3B), including several proteases and anti-proteases (ADAMTSs & SERPINs), elastogenesis genes (*ELN, EMILIN1, FBLN1, FBLN5 & MFAP4*), and collagens (*COL6A1, COL6A2 & COL18A1*).

**Figure 3:**
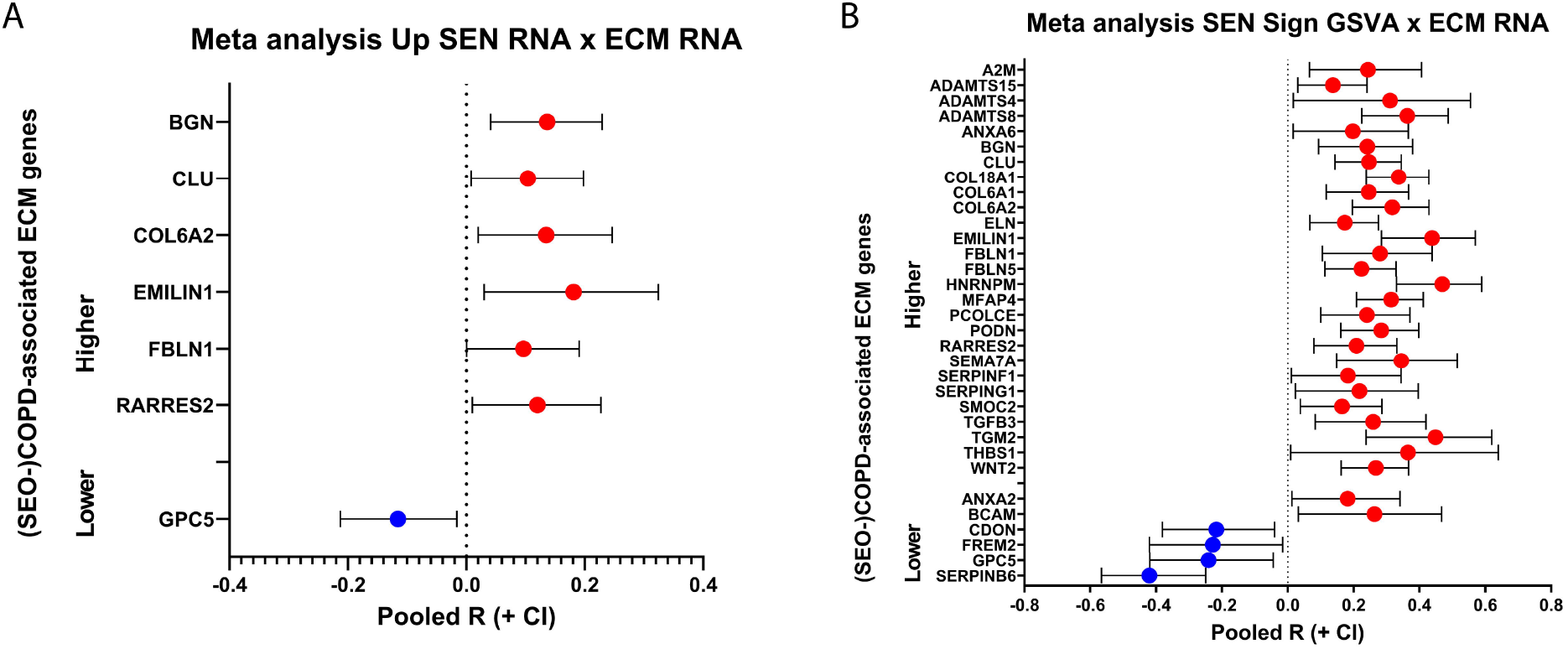
Consistent associations between (SEO-)COPD-associated ECM genes and senescence genes and signatures. Meta analyses were performed on the correlation results of the five up-regulated senescence genes (A) and the four senescence signatures (B) with (SEO-)COPD-associated ECM genes in lung tissue. Significant ECM (P<0.05) are depicted in the forest plots with positive associations in red and negative associations in blue. The pooled R with confidence intervals are shown on the x-axis.

### In situ verification of senescence association with FBLN5 and COL6 in lung parenchyma and small airway walls

Since the tissue homogenates used for the previous analyses are a mix of peripheral lung regions, we aimed to verify our findings in situ using histology assessing lung parenchyma and small airway walls separately ^(17)^. In the parenchymal region, positive correlations were observed for COL6A1 staining intensity and FBLN5 positively stained area with p21 positively stained area (Figure 4A+C), with similar trends towards significance for COL6A1 area and FBLN5 intensity (Figure 4A). In the small airway walls, positive correlations were observed for COL6A1 area, COL6A2 intensity, and FBLN5 area and intensity with p21 area (Figure 4B). So, we confirmed the association between p21 and the ECM proteins COL6A1, COL6A2, and FBLN5 both in parenchymal lung regions and small airway walls, with the strongest associations for COL6A1.

**Figure 4:**
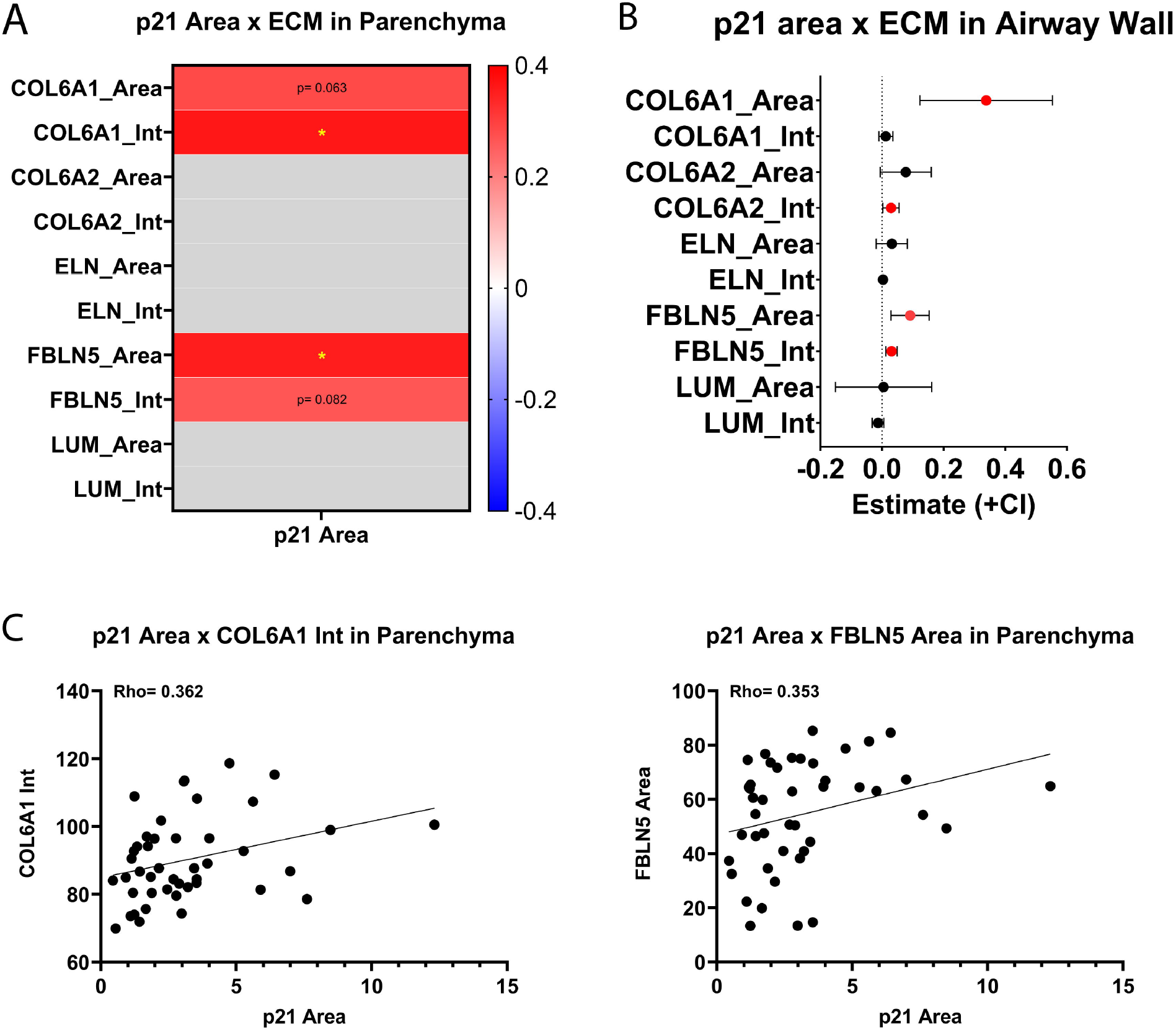
Verification of correlations between p21 and ECM in situ. Data from IHC stainings for p21 and ECM on COPD and non-COPD lung tissue was used to verify the found ECM-senescence in situ. Percentage area of p21 positive staining (p21 area) was correlated with ECM area or intensity of ECM staining (Int) in the parenchyma (A) using Spearman’s correlations (A) or in the airway walls (B) using a linear mixed model (LMM) to correct for multiple airways per patient. Pearson’s correlation coefficient (Rho) of (trend towards) significant correlations are shown in the heatmap (A). P-value is indicated for a trend (0.05 < P < 0.1) and an asterisk indicates significant correlations (P<0.05). The LMM estimates and confidence intervals for the correlations in the airway walls are depicted in the forest plot and colored when significant (P<0.05). The significant correlations in the parenchyma are depicted in the scatter plots (C) with the Spearman’s correlation coefficients (rho) in the top left corner of the graphs.

### Confirmation of senescence-associated ECM-related genes in primary lung fibroblasts identified consistent associations for proteases, elastogenesis genes, and COL6

Since fibroblasts are the main producers of the ECM and given our previous observations in lung fibroblasts, we decided to confirm the ECM-senescence correlations identified *in vivo* in primary lung fibroblasts. While in lung tissue most ECM genes were correlated with *CDKN1A/p21*, in fibroblasts most ECM genes (11 genes) were correlated with *CDKN2A/p16* (Figure 5A). Ten ECM genes were correlated with *LMNB1* and 6 ECM genes correlated with *CDKN1A/p21*, with only 3 in the same direction as observed in lung tissue. The meta-analysis in lung fibroblasts revealed 14 ECM genes to be consistently associated with the up-regulated senescence genes and 4 with similar trends (Figure 5C), of which 14 in the same direction as observed in lung tissue, including some proteases *(ADAMTS1 & ADAMTS15)* and anti-proteases (*SERPINE2 & SERPING1*), elastogenesis genes (*ELN, EMILIN1 & FBLN1*), and *COL6A1* and *COL6A2*.

**Figure 5:**
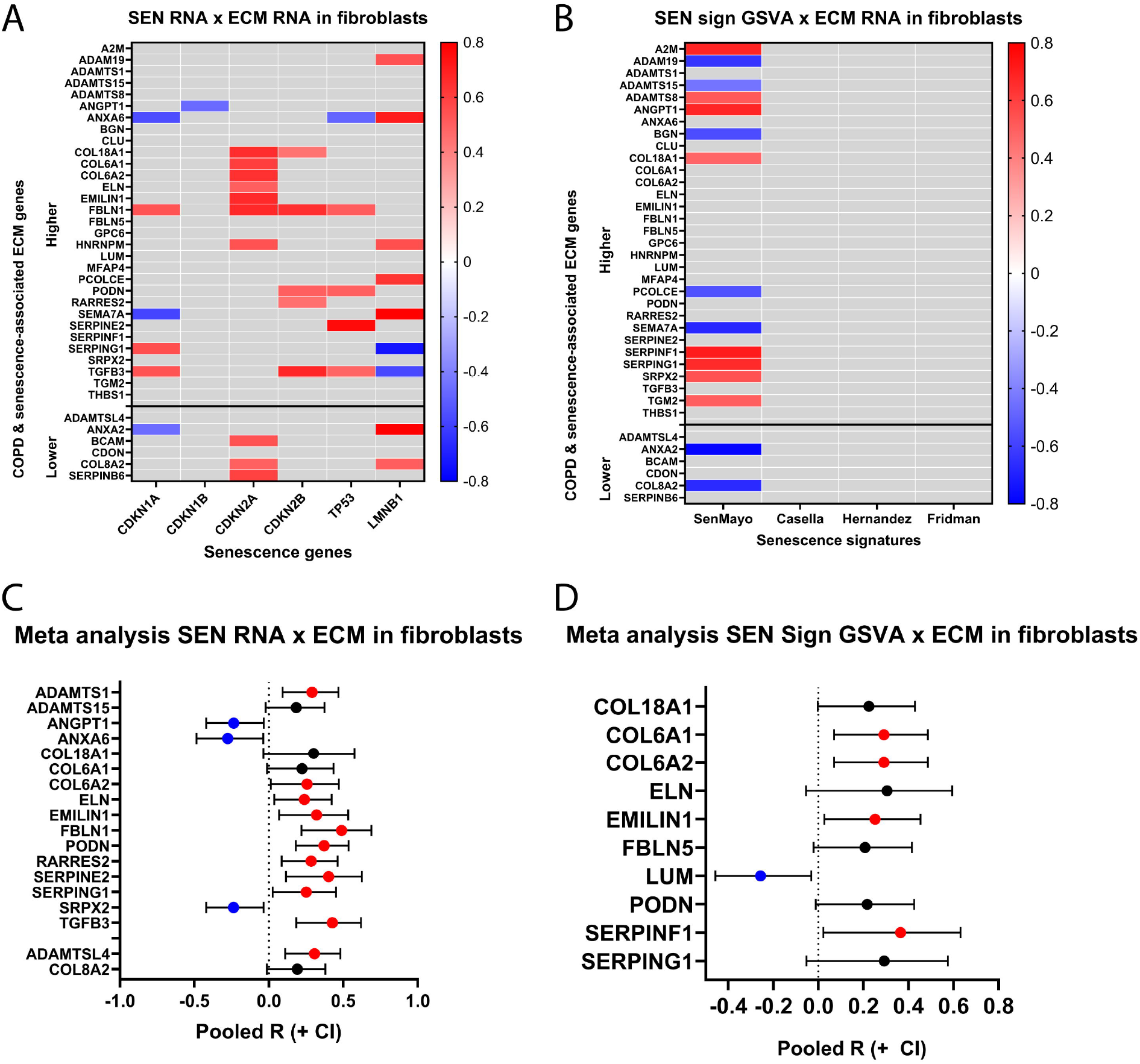
Confirmation of ECM-senescence correlations in primary human lung fibroblasts. Gene expression was measured in primary human lung fibroblasts from COPD (n=10) and non-COPD patients (n=11) at baseline culture using RNA sequencing. COPD- and senescence-associated ECM-related genes in lung tissue were correlated with senescence gene expression (A) using Pearson’s correlations and with the calculated GSVA scores of the senescence signatures (B) using Spearman’s correlations and depicted in the heatmap (A). Heatmaps depict the correlations coefficients (r or rho) for significant correlations (P<0.05) with positive correlations in red and negative correlations in blue. The ECM-related genes and proteins higher expressed in COPD are depicted on top and the ones lower expressed in COPD are depicted on the bottom of the y-axis. The senescence genes and GSVA scores of senescence signatures are depicted on the X-axis. Meta analyses were performed using the correlation results for the senescence genes (C) and signatures (D) and the pooled R and confidence intervals are depicted in the forest plots when a trend towards significance was observed. Significant ECM in the meta-analyses are colored.

Next, the COPD- and senescence-associated ECM genes were correlated with the senescence signature scores in fibroblasts. Significant correlations were only observed for the SenMayo score, with 15 significant ECM genes, of which 8 genes were correlated in the same direction as observed in lung tissue. A meta-analysis was performed to assess the consistency of the ECM-senescence associations across the different senescence signatures, which resulted in 5 significant ECM genes and 5 with similar trends (Figure 5D), of which 9 ECM genes being consistently associated in the same direction as observed in lung tissue, including *COL6A1* and *COL6A2* and the elastogenesis genes (*ELN, EMILIN1 & FBLN5*).

### Functional validation in senescence-induced parenchymal lung fibroblasts

To investigate a potential causal effect of senescence on the ECM changes, we assessed the senescence-associated ECM genes identified in lung tissue in our PQ-induced senescence model ^(12)^ (Supplemental figure E2). Three ECM genes, *SERPINE2, GPC6*, and *COL8A2*, were functionally validated in the same direction as observed in lung tissue with a similar trend for *ADAMTS1* (Figure 6A). However, most ECM genes were down-regulated at the transcript level in PQ-induced senescent fibroblasts, which is in opposite direction as its association in lung tissue. Together with our previous observations in lung fibroblasts, where we showed increased SASP protein secretion and lower ECM expression upon PQ-induced senescence ^(12, 19)^, it is likely that the senescence-induced fibroblasts demonstrate a phenotype switch towards high SASP secretion and low ECM production, at least *in vitro*. To address this further, we investigated gene expression of the SenMayo signature, which includes majorly SASP genes, in our PQ-induced senescence model and observed an up-regulation of many (27 genes) of these genes, with the strongest increase of well-known SASP genes *CXCL8, MMP3*, and *GDF-15* (Figure 6B), as also observed at secretion level in our previous study ^(19)^.

**Figure 6:**
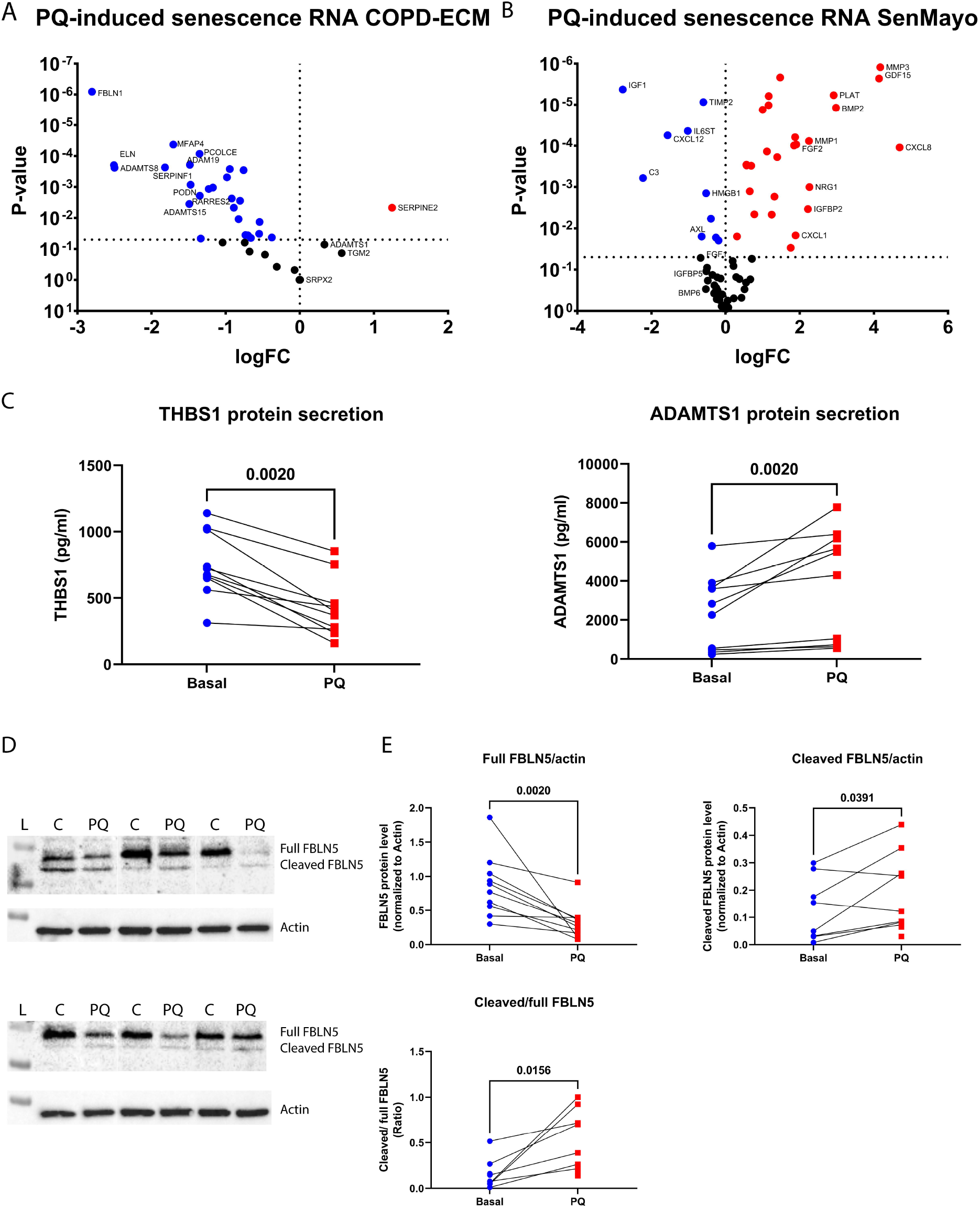
Functional validation of ECM-senescence correlations in senescence-induced primary human lung fibroblasts. Senescence was induced using 250μM Paraquat in primary human lung fibroblasts derived from non-COPD controls (n=11). Gene expression was measured using RNA sequencing, secreted protein levels were measured using ELISA, and FBLN5 protein levels were measured using Western Blot. Gene expression of COPD- and senescence-associated ECM (A) and genes from the SenMayo signature (B) were compared between Paraquat-induced and untreated fibroblasts using generalized linear models. Logarithmic fold changes (x-axis) and p-values (y-axis) are depicted in the volcano plots and colored when significant (P<0.05). Secreted protein levels of THBS1 and ADAMTS1 are shown in the dot plots (C). Examples of representative Western Blot membranes with visualization of full FBLN5, cleaved FBLN5 and Actin are shown (D). L = protein ladder, C = untreated fibroblasts & PQ = PQ-induced senescent fibroblasts. Quantification of total intensity of Full size FBLN5, Cleaved FBLN5 and the ratio of Cleaved FBLN5/Full size FBLN5 of untreated and PQ-induced senescent fibroblasts are shown in the dot plots (E). The lines represent matching donors. Technical outliers were excluded. Statistical significance was tested using Wilcoxon signed-rank test, p-values are indicated and bold when < 0.05.

In addition, we assessed the secretion of the three strongest senescence-associated ECM observed in lung tissue in our PQ-induced senescent fibroblasts. ADAMTS1 protein secretion was increased after senescence induction, while, on the other hand, protein secretion of THBS1 was decreased (Figure 6C). The levels of secreted ADAMTS4 were below detection limit.

Since FBLN5 was positively correlated with senescence on protein level and confirmed with histology, we also assessed FBLN5 protein levels in our senescence induction model. Of interest, proteolytic cleavage of FBLN5 by serin proteases has been described to result in a 10 kDa shorter none-functional cleaved protein variant ^(20)^, which was found in higher levels in COPD lung tissue before ^(21)^. After senescence induction the protein levels of the full length FBLN5 protein (± 50 kDa) were decreased (Figure 6D+E). However, the absolute levels of cleaved FBLN5 (± 40 kDa) and the ratio of cleaved FBLN5 over full length FBLN5 increased after senescence induction (Figure 6E), suggesting more cleavage and non-functional FBLN5 protein with senescence.

Please find a summarizing overview of most consistent and confirmed ECM-senescence correlations in situ and in primary fibroblasts in table 2.

**Table 2:**
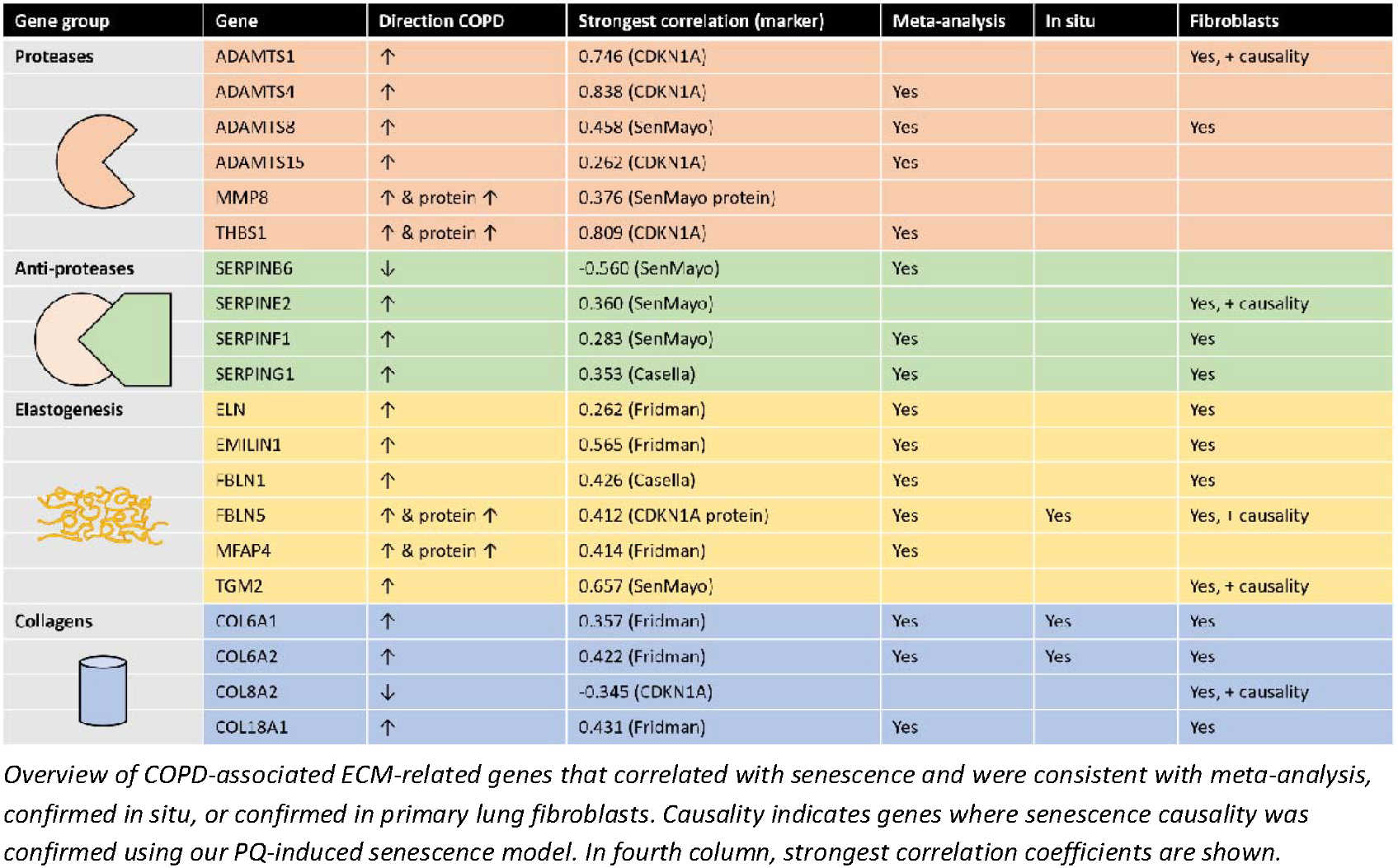
Summary overview of consistent and confirmed ECM-senescence correlations.

## DISCUSSION

This study aimed to determine whether COPD-associated ECM changes correlate with cellular senescence in lung tissue and whether the previously observed ECM dysregulation in senescent lung fibroblasts translates *in vivo* contributing to COPD and SEO-COPD pathology. Indeed, we confirmed that many COPD-, and in particular SEO-COPD-, associated ECM changes are correlated with senescence in lung tissue, especially with *CDKN1A/p21* and the SenMayo and Fridman senescence signature scores. The strongest correlations include proteases, regulators of ECM organization, and the most consistent correlations include the elastogenesis genes and 2 genes encoding collagen 6, *COL6A1* and *COL6A2* (See Table 2 for an overview). The associations between senescence and *COL6A1, COL6A2*, and *FBLN5* were confirmed with histology both in lung parenchyma and small airway walls and in primary lung fibroblasts. In addition, we confirmed senescence correlations for some proteases and anti-proteases, elastogenesis genes in primary lung fibroblasts. Finally, we showed that cellular senescence may play a role in the cleavage of FBLN5, resulting in a non-functional protein, which was found in higher levels in COPD lung tissue before.

The strongest correlations between ECM and senescence in lung tissue were observed for proteases, including *THBS1, ADAMTS4*, and *ADAMTS1*. Some of these proteases were consistently associated with the senescence scores (*ADAMTS4, ADAMTS8, ADAMTS15*, and *THBS1*) and confirmed in primary lung fibroblasts (*ADAMTS1, ADAMTS8*, and *ADAMTS15*). In addition, ADAMTS1 secretion was functionally validated after PQ-induced senescence. These results indicate that senescence may affect the protease-antiprotease balance. While the protease-antiprotease imbalance has long been recognized to play a role in the pathogenesis of COPD ^(22)^, this was mainly attributed to immune cells, whereas our results indicate also involvement of fibroblast senescence. Interestingly, in other tissues it has already been demonstrated that senescent cells have higher expression and secretion of proteases, including MMP and ADAM proteins ^(4, 23, 24)^. In line with this, in a previous study we found higher secretion levels of multiple MMP proteins upon PQ-induced senescence in lung fibroblasts ^(19)^, including MMP-9 (reported), and MMP-2, -3, and -10 (data not shown). Together, these data support the hypothesis that cellular senescence, including fibroblast senescence, may contribute to dysregulated ECM homeostasis in COPD through an imbalance in proteases and antiproteases. Future studies should further explore the role of fibroblast senescence in this imbalance, including measuring protease activity.

Next to the strong association for proteases, consistent ECM-senescence associations were observed for the elastogenesis genes, of which many have been linked to COPD before ^(21, 25-29)^. Our group has described the role of elastogenesis genes in COPD before and proposed that these genes were up-regulated as a dysfunctional repair response ^(21)^. We now demonstrate the association between these elastogenesis genes (*ELN, FBLN1, FBLN5, EMILIN1, MFAP4*, and *TGM2*) and senescence. In skin, epidermal senescence resulted in altered elastic fiber morphology and structure ^(30)^, but the exact role of senescence in disturbed elastic fiber formation in lungs remains unclear. Most elastogenesis genes were confirmed in primary lung fibroblasts, but no causal effect was found in PQ-induced senescent fibroblasts, indicating that up-regulation of these genes may be an indirect effect of senescence. We hypothesize that in a senescence rich environment more proteases are produced resulting in active breakdown of ECM and elastic fibers. This in turn will lead to an increased need for repair and thus increased expression of elastogenesis genes. This combination of senescence-induced proteolysis and insufficient repair may contribute to emphysema in COPD patients.

The senescence association of FBLN5 protein level was also confirmed with histology and more cleavage of FBLN5 was found in senescence-induced fibroblasts. FBLN5 has been demonstrated to be a key player in elastic fiber formation and when FBLN5 is knocked out in mice it disturbs elastic fiber formation with these mice developing emphysema ^(31)^. Cleavage of FBLN5 leads to a non-functional protein and *in vitro* this leads to disturbed elastogenesis ^(31)^. The level of this non-functional cleaved FBLN5 was observed to be higher in COPD lung tissue ^(21)^. Here, we demonstrated that senescent lung fibroblasts can contribute to these higher levels of non-functional cleaved FBLN5 in COPD lung tissue, which is associated with changes in the other elastogenesis genes as well, further adding to the insufficient repair in COPD.

The strong and consistent positive correlations between senescence and collagen 6 were confirmed with histology and in primary lung fibroblasts, but expression of both *COL6A1* and *COL6A2* was down-regulated in senescence-induced fibroblasts. In addition, our group previously demonstrated that both collagen 6 proteins in lung tissue were associated with age ^(32)^. These results may indicate that in a senescence rich environment *in vivo* collagen 6 may be induced via an indirect effect of senescence. This indirect effect may be a repair response to protease-induced tissue damage or by activating TGF signaling. In addition, the pro-inflammatory and proteolytic local environment in the COPD lungs, partly driven by SASP from senescent fibroblasts, may also induce TGF signaling and subsequently COL6A1 and COL6A2 production ^(33, 34)^. In line with this hypothesis, multiple activators of TGF signaling were associated with senescence in lung tissue, including *BGN, TGFB3, TGM2*, and *THBS1*. In accordance with our previous study ^(12)^, we found a down-regulation of structural ECM genes and an up-regulation of pro-inflammatory SASP proteins in PQ-induced senescent fibroblasts. We hypothesize that these senescent fibroblasts *in vitro* have a shifted phenotype from ECM production and regulation towards a pro-inflammatory SASP phenotype that contributes to ECM dysregulation in COPD lungs. In senescent dermal fibroblasts this shift in phenotype was indeed observed before with reduced synthesis of structural ECM proteins, including COL1A1, COL3A1, and COL4A1 ^(35, 36)^. However, no other studies have demonstrated this phenotype shift in senescent lung fibroblasts yet, and it remains unclear how this translates to the *in vivo* situation. Spatial techniques are needed to confirm this phenotype shift *in vivo* in COPD lungs.

The limitation of a cross-sectional study like ours is that changes in structural ECM proteins that may have accumulated over a long time may not be easily detected *in vivo* and may also follow different timelines compared to our, more acute, *in vitro* model. Due to this mismatch, causal validation in our PQ-induced senescence model in fibroblasts turned out to be challenging. Another factor to consider here is that different senescence inducers lead to different senescent phenotypes, which may differ in disease context as well. We used our Paraquat model as this induces senescence by an increase in oxidative stress, likely the main contributor to senescence in COPD ^(37-39)^, but other senescence induction models could be considered as well. Nevertheless, we successfully confirmed causal effects of senescence induction for four genes, including for protein secretion *ADAMTS1*. Furthermore, the observed associations *in vivo* can result from different cell types and different lung regions and can also include indirect effects. To account for this, we confirmed our findings with histology and in primary lung fibroblasts, both resulting in consistent associations for collagen 6 and FBLN5. The importance of indirect effects is supported by the strong senescence associations with proteases and the SASP in the SenMayo signature. Finally, in our *in vitro* model we only assessed the effect of senescence on ECM changes, but it should be kept in mind that altered ECM composition, structure and organization may also influence cellular senescence, as reviewed by Blokland et al ^(40)^.

In conclusion, after demonstrating a link between ECM dysregulation and senescence in COPD-derived lung fibroblasts previously, we have now demonstrated that many (SEO-)COPD-associated ECM-related genes and proteins are correlated with senescence *in vivo* in COPD lung tissue. As the strongest correlations include proteases and elastogenesis genes, our findings suggest that senescence can contribute to ECM dysregulation in COPD by affecting the protease-antiprotease imbalance and the disturbance of elastic fiber formation, with likely a contributing role for senescent fibroblasts. The exact mechanism of how senescent cells affect elastic fiber regulation in COPD lungs should be investigated in more detail, as well as the potential indirect and chronic effects of senescence on ECM dysregulation. Combined longitudinal and spatial approaches are needed, connected to more complex *in vitro* and *ex-vivo* models. Altogether, this will lead to a better understanding of the functional consequences of increased senescence on disease pathology in COPD patients and may open new opportunities to develop therapeutic strategies to restore ECM regulation in COPD by targeting senescence.

## Supporting information

Online suppelment

