## Supplementary material for "Cellular Senescence Affects ECM Regulation in COPD Lung Tissue": Online suppelment

**ONLINE SUPPLEMENT**

**FULL DESCRIPTION OF MATERIALS AND METHODS**

**Lung tissue collection for transcriptomic and proteomic analyses**

For the PRoteogenomics Early onSeT cOpd (PRESTO) lung cohort, peripheral lung tissue was collected from never, ex-, and current smoking non-COPD controls (FEV_1_/FVC > 70% of predicted), from ex- and current smoking COPD patients (FEV_1_/FVC < 70% of predicted) with GOLD stage II, III, and IV, and from COPD patients with alpha-1 antitrypsin deficiency (n=117 subjects total). These lung tissues were obtained from left-over material from lung transplantation or tumor resection surgery. Lung tissue from tumor resection materials was taken far away from the tumor and microscopically checked for abnormalities by a lung pathologist. For the current study, COPD patients with alpha-1 antitrypsin deficiency and never smoker controls were excluded.

The study protocol was consistent with the Research Code of the University Medical Center Groningen (research code UMCG (umcgresearch.org) and national ethical and professional guidelines (“Code of conduct for Health Research; coreon.org). Lung tissues used in this study were derived from leftover lung material after lung surgery from archival materials and were exempt from consent in compliance with applicable laws and regulations (Dutch laws: Medical Treatment Agreement Act (WGBO) art 458 / GDPR art 9/ UAVG art 24). This material was not subject to the Medical Research Human Subjects Act in the Netherlands, confirmed by a statement from the University Medical Center Groningen. All samples and clinical information were coded before experiments were performed, blinding all directly identifying information to the investigators.

**Lung tissue transcriptomics**

Transcriptomic data from 60 COPD patients and 30 non-COPD controls was used. Briefly, lung tissue cryo sections were used for RNA isolation using the AllPrep DNA/RNA/miRNA Universal Kit (Qiagen) according to manufacturer’s protocol. RNA sequencing was performed by GenomeScan B.V. on an Illumina Novaseq 6000 (PE 2x150bp). Raw RNA sequencing reads were aligned to GRCh38 reference (Ensembl gene build 100) using STAR aligner (ver. 2.7.3a) using multi-sample 2-pass mapping. Equivalent alignments with duplicate UMIs were discarded. Read counts per gene were quantified with HtSeq package (ver. 0.12.4). Differential expression analyses were performed using edgeR (ver3.40.2) depending on Limma (ver. 3.54.2) in R (ver. 4.2.2). A list of 471 ECM(-related) genes based on three Gene Ontology databases (GO:0085029, GO:0062023, GO:0005201) was used for differential expression analyses. ECM(-related) gene expression was compared between control current smokers (CS) and control ex-smokers (ES), between COPD and control, and between SEO-COPD and control (see table 1 for characteristics) using generalized linear models corrected for age and sex. The (SEO-)COPD-associated ECM(-related) genes were correlated with six major senescence genes (*CDKN2A, CDKN1A, TP53, CDKN2B, CDKN1B,* and *LMNB1*) using Pearson’s correlations in all available lung tissue samples. GSVA scores of four senescence signatures (SenMayo ^(1)^, Casella ^(2)^, Hernandez-Segura ^(3)^, and Fridman ^(4)^) were calculated using the GSVA package (ver. 1.50.5) with exclusion of the (SEO-)COPD-associated ECM(-related) genes. The (SEO-)COPD-associated ECM(-related) genes were correlated with these GSVA scores using Spearman’s correlations as these were not normally distributed. Multiple testing correction was done using the Benjamini-Hochberg Procedure. A false discovery rate (FDR) below 0.05 was considered statistically significant. Meta analyses were performed on the correlation results between ECM genes and senescence genes and senescence signatures using metafor package (ver. 4.8-0). P-value below 0.05 was considered statistically significant.

**Lung tissue proteomics**

From the same lung tissue cryo samples as used for transcriptomics, serial sections were used for proteomics. Total protein lysates were prepared using RIPA lysis buffer, and the proteins were digested into peptides using in-gel digestion as described previously for other tissues ^(5)^. Discovery-based proteomics protein identifications were done using the previously described data independent acquisition method on an Orbitrap 480 LC-MS/MS platform ^(6)^. LC-MS raw data were processed with Spectronaut (ver. 14.9.201124, Biognosys) using the standard settings of the directDIA workflow with a human SwissProt database ([www.uniprot.org](http://www.uniprot.org), 20350 entries). For the quantification, local normalization based on locally weighted regression and smoothing scatter plot (LOWESS) was applied and the Q-value filtering was adjusted to the 0.2 percentile. The numerical output of the protein peak areas was log transformed using R and missing values were imputed with a fixed value of 0.5. Differential expression analyses were performed using edgeR (ver. 3.40.2) depending on Limma (ver. 3.54.2) in R (ver. 4.2.2). Protein levels of the ECM(-related) proteins (as described above) were compared between control current smokers (CS) and control ex-smokers (ES), between COPD and control, and between SEO-COPD and control (see table 1 for characteristics) using generalized linear model corrected for age and sex. The (SEO-)COPD-associated ECM(-related) proteins were correlated with the six major senescence genes using Pearson’s correlations and with these GSVA scores using Spearman’s correlations in all available lung tissue samples. Multiple testing correction was done using the Benjamini-Hochberg Procedure. A false discovery rate (FDR) below 0.05 was considered statistically significant.

**Immunohistochemical confirmation on lung tissue**

Data from the HOLLAND study regarding COL6A1, COL6A2, ELN, FBLN5, and LUM staining in lung tissue, described by Joglekar et al. ^(7)^ and p21 staining, described by Chen et al. ^(8)^ was used to verify ECM-senescence correlations in situ using lung tissue sections from ex-smoking COPD patients (n=27) and ex-smoking non-COPD controls (n=18) to avoid smoke effects. Representative images of IHC staining can be found in the original studies. Area of positive p21 staining was correlated with area of positive ECM staining and intensity of ECM staining in the parenchymal and airway wall regions for the available COPD- and senescence-associated ECM: COL6A1, COL6A2, ELN, FBLN5, and LUM. For the parenchymal regions, Spearman’s correlations were performed as data was not normally distributed and in the airway walls, Linear Mixed Models were performed to correlate per airway and to correct for multiple airways per patient. For verification, P-value below 0.05 was considered statistically significant.

**Senescence-induced lung fibroblast experiments**

Primary human lung fibroblasts from SEO-COPD (n=10) and non-COPD patients (n=11) (see Table E1 for patient characteristics) were cultured at baseline conditions and cellular senescence was induced in non-COPD-derived fibroblasts using our previously described Paraquat-induced senescence model (Woldhuis et al. AJP 2020) and RNA sequencing was performed. Briefly, senescence was induced using 250uM Paraquat in T25 flasks and 5 days after Paraquat treatment the fibroblasts were reseeded into 24-wells plates (75,000 cells per well) at ± 90% confluency. After three days, RNA using Trizol (Invitrogen) was collected and proteins using RIPA lysis buffer (Thermo Scientific) were collected from the wells. Senescence induction was confirmed by SA-β-gal staining, reduced proliferation in T25 flasks, and *CDKN1A* (p21) and *LMNB1* gene expression, as described before ^(9)^. RNA was isolated using the AllPrep DNA/RNA/miRNA Universal Kit (Qiagen) according to manufacturer’s protocol. RNA sequencing was performed by GenomeScan B.V. on an Illumina Novaseq 6000 v1.5. Raw RNA sequencing reads were aligned to GRCh38 reference (Ensembl gene build 100) using STAR aligner (ver. 2.7.3a) using multi-sample 2-pass mapping. Equivalent alignments with duplicate UMIs were discarded. Read counts per gene were quantified with HtSeq package (ver. 0.12.4). Differential and expression and correlation analyses were performed using edgeR (ver. 3.42.4) depending on Limma (ver. 3.56.2) in R (ver. 4.3.1). The (SEO-)COPD- and senescence-associated ECM(-related) genes were correlated with the six senescence genes using Pearson’s correlations. GSVA scores of the four senescence signatures were calculated using the GSVA package (ver. 1.50.5) with exclusion of the (SEO-)COPD-associated ECM(-related) genes. The (SEO-)COPD-associated ECM(-related) genes were correlated with these GSVA scores using Spearman’s correlations as these were not normally distributed. Meta analyses were performed on the correlation results between ECM genes and senescence genes and senescence signatures using metafor package (ver. 4.8-0). For functional validation, P-value below 0.05 was considered statistically significant.

Next, expression of the (SEO-)COPD- and senescence-associated ECM(-related) genes were compared between PQ-induced and untreated fibroblasts. Finally, to assess SASP gene expression, expression of genes from the SenMayo signature were compared between PQ-induced and untreated fibroblasts. For functional validation, P-value below 0.05 was considered statistically significant.

**Secreted protein analyses**

Cell-free supernatants were collected and stored at -80°C prior to ELISA analyses. Secreted ADAMTS1, ADAMTS4, and THBS1 were measured using Human DuoSet ELISA (R&D Systems).

**Western Blot fibroblast samples**

To assess FBLN5 protein levels, Western Blot was performed on collected protein samples from untreated and senescence-induced fibroblasts. Protein samples were centrifuged at 4°C for 10 minutes at maximum speed. Laemmli sample buffer was added to the protein samples, and these were separated on an SDS-PAGE gel and blotted on a nitrocellulose membrane (Bio-Rad Laboratories, Lunteren, The Netherlands). Membranes were blocked with 5% fat free milk in TBST and incubated with primary antibody against FBLN5 (Novus) in 5% fat free milk in TBST overnight at 4°C. The next day, membranes were incubated with the secondary antibody Goat anti-mouse HRP (DAKO) in 5% fat free milk in TBST for one hour at RT. Antibody binding was visualized with SuperSignal West Pico PLUS Chemiluminescent Substrate (Thermo Scientific) using a ChemiDoc XRS+ imager (Bio-Rad Laboratories). To quantify protein levels, Image Lab Software (Bio-Rad Laboratories) was used for densitometry of detected protein bands and actin was used as loading control.

**SUPPLEMENTARY TABLES AND FIGURES**

**Table E1: Patient characteristics of primary lung fibroblasts derived from SEO-COPD and non-COPD patient.**

| **Variable** | **Non-COPD ex-smoker control** | **SEO-COPD patients** |
| --- | --- | --- |
| **Number** | 10 | 11 |
| **Age, median (range)** | 55 (36-60) | 51 (44-55) |
| **Male/female, N** | 3/7 | 3/8 |
| **Pack-years, median (range)** | 25 (10-40) | 27 (6-54) |
| **FEV_1_ %pred, median (range)** | 89.1 (70.0 – 117.1) | 17.1 (12.0 – 28.3) |
| **FEV_1_/FVC, median (range)** | 75.6 (70.0 – 86.6) | 26.7 (22.0 – 45.5) |

**Table E2: Results of transcriptomics analysis of COPD-associated ECM genes**

| **Gene** | **LogFC** | **P-Value** | **P adjust** |
| --- | --- | --- | --- |
| **VWA2** | -0.673204 | 2.66E-07 | 0.000090 |
| **ADAMTS15** | 0.712327 | 0.000043 | 0.005471 |
| **SERPING1** | 0.360423 | 0.000049 | 0.005471 |
| **FBLN5** | 0.443042 | 0.000205 | 0.012739 |
| **CXCL12** | 0.600417 | 0.000292 | 0.012739 |
| **SRPX2** | 0.461534 | 0.000295 | 0.012739 |
| **PODN** | 0.553314 | 0.000295 | 0.012739 |
| **SERPINB6** | -0.222378 | 0.000306 | 0.012739 |
| **HAS3** | -0.702627 | 0.000339 | 0.012739 |
| **SMOC2** | 0.591702 | 0.000601 | 0.020302 |
| **FREM2** | -0.466403 | 0.001053 | 0.032371 |
| **ANOS1** | -0.390363 | 0.001207 | 0.033990 |

**Table E3: Results of proteomics analysis of COPD-associated ECM proteins**

| **Protein** | **LogFC** | **P-value** | **P adjust** |
| --- | --- | --- | --- |
| **FBLN5** | 0.155706 | 0.000042 | 0.007364 |
| **CTSH** | -0.172830 | 0.000180 | 0.014460 |
| **MMP8** | 1.293908 | 0.000246 | 0.014460 |
| **ANXA6** | 0.055977 | 0.000727 | 0.031969 |

**Table E4: Results of transcriptomics analysis of SEO-COPD-associated ECM genes**

| **Gene** | **LogFC** | **P-Value** | **P adjust** |
| --- | --- | --- | --- |
| **CXCL12** | 1.135935 | 1.96E-06 | 0.000657 |
| **ADAMTS15** | 1.213344 | 4.46E-06 | 0.000747 |
| **PODN** | 0.949707 | 7.00E-06 | 0.000768 |
| **VWA2** | -0.883368 | 9.17E-06 | 0.000768 |
| **SMOC2** | 1.084586 | 0.000027 | 0.001780 |
| **CDON** | -0.756654 | 0.000058 | 0.003234 |
| **EMILIN1** | 0.877560 | 0.000112 | 0.005376 |
| **FBN1** | 0.618627 | 0.000210 | 0.008811 |
| **SERPING1** | 0.550754 | 0.000292 | 0.009795 |
| **FBLN1** | 0.730349 | 0.000292 | 0.009795 |
| **SERPINB6** | -0.362655 | 0.000333 | 0.010148 |
| **SERPINF1** | 0.772836 | 0.000366 | 0.010221 |
| **ANGPT1** | 0.664065 | 0.000535 | 0.012020 |
| **S100A10** | -0.581744 | 0.000536 | 0.012020 |
| **ANXA2** | -0.387563 | 0.000538 | 0.012020 |
| **ANXA6** | 0.451869 | 0.000611 | 0.012801 |
| **COL6A1** | 0.531120 | 0.000760 | 0.014474 |
| **COL8A2** | -0.654763 | 0.000778 | 0.014474 |
| **SRPX2** | 0.736618 | 0.000826 | 0.014558 |
| **CILP** | 1.237788 | 0.000943 | 0.014571 |
| **ADAMTSL4** | -0.581424 | 0.000954 | 0.014571 |
| **TGM2** | 0.599495 | 0.001017 | 0.014571 |
| **FBLN5** | 0.627746 | 0.001028 | 0.014571 |
| **HAS3** | -1.034874 | 0.001049 | 0.014571 |
| **RARRES2** | 0.612844 | 0.001087 | 0.014571 |
| **ANOS1** | -0.663434 | 0.001218 | 0.015698 |
| **A2M** | 0.548779 | 0.001544 | 0.019151 |
| **THBS1** | 1.739028 | 0.001911 | 0.022859 |
| **COL6A2** | 0.479296 | 0.001991 | 0.023004 |
| **COL18A1** | 0.505935 | 0.002070 | 0.023110 |
| **COL14A1** | 0.806370 | 0.002143 | 0.023157 |
| **SEMA7A** | 0.684255 | 0.002256 | 0.023612 |
| **ACAN** | 1.997186 | 0.002476 | 0.025139 |
| **NID2** | 0.457887 | 0.002674 | 0.026279 |
| **LAMB1** | 0.571150 | 0.002746 | 0.026279 |
| **PCOLCE** | 0.655602 | 0.002842 | 0.026449 |
| **TGFB3** | 0.684961 | 0.003293 | 0.029813 |
| **LTBP1** | 0.536120 | 0.003596 | 0.031034 |
| **ADAM19** | 0.635483 | 0.003613 | 0.031034 |
| **LUM** | 0.701719 | 0.003870 | 0.032413 |
| **ELN** | 0.974050 | 0.004342 | 0.035478 |
| **WNT2** | 0.668594 | 0.004765 | 0.037168 |
| **MFAP4** | 0.636708 | 0.004937 | 0.037168 |
| **SERPINE2** | 0.841291 | 0.004956 | 0.037168 |
| **PRG4** | 1.512691 | 0.004993 | 0.037168 |
| **ADAMTS4** | 2.325955 | 0.005617 | 0.040751 |
| **FREM2** | -0.635069 | 0.005717 | 0.040751 |
| **ADAMTS8** | 0.713677 | 0.006050 | 0.042227 |
| **ANGPT4** | -1.278532 | 0.006599 | 0.045115 |
| **CLU** | 0.492198 | 0.006886 | 0.045288 |
| **ADAMTS1** | 0.987691 | 0.006895 | 0.045288 |
| **GPC6** | 0.762153 | 0.007121 | 0.045874 |
| **GPC5** | -0.770131 | 0.007452 | 0.047100 |
| **BGN** | 0.496298 | 0.007915 | 0.048316 |
| **BCAM** | -0.364890 | 0.007933 | 0.048316 |
| **COL6A5** | 0.778835 | 0.008088 | 0.048383 |
| **HNRNPM** | 0.262556 | 0.008271 | 0.048608 |

**Table E5: Results of proteomics analysis of SEO-COPD-associated ECM proteins**

| **Protein** | **LogFC** | **P-value** | **P adjust** |
| --- | --- | --- | --- |
| **FBLN5** | 0.265841 | 0.000034 | 0.005924 |
| **CTSH** | -0.287127 | 0.000203 | 0.017843 |
| **TINAGL1** | 0.431378 | 0.000385 | 0.022576 |
| **PSAP** | -0.454147 | 0.000976 | 0.034242 |
| **LGALS3BP** | -0.086342 | 0.001038 | 0.034242 |
| **TGM2** | 0.327208 | 0.001328 | 0.034242 |
| **THBS1** | 0.300548 | 0.001362 | 0.034242 |
| **MFAP4** | 0.274556 | 0.001620 | 0.035636 |
| **CLU** | 0.201646 | 0.002181 | 0.042643 |

**Table E6: Results of correlations between senescence genes and (SEO-)COPD-associated ECM genes**

| **R values** | **CDKN1A** | **CDKN1B** | **CDKN2A** | **CDKN2B** | **TP53** | **LMNB1** |
| --- | --- | --- | --- | --- | --- | --- |
| **A2M** | 0.126 | 0.090 | -0.174 | -0.004 | 0.201 | 0.153 |
| **ACAN** | 0.004 | 0.025 | 0.032 | -0.173 | 0.148 | 0.176 |
| **ADAM19** | **0.541** | -0.266 | 0.196 | -0.086 | -0.011 | **0.717** |
| **ADAMTS1** | **0.746** | -0.280 | -0.139 | -0.095 | -0.267 | **0.298** |
| **ADAMTS15** | **0.262** | 0.001 | -0.045 | -0.142 | -0.048 | 0.024 |
| **ADAMTS4** | **0.838** | **-0.524** | 0.079 | -0.111 | -0.152 | **0.542** |
| **ADAMTS8** | **0.429** | **-0.325** | 0.014 | 0.205 | 0.081 | 0.200 |
| **ANGPT1** | -0.024 | 0.266 | -0.338 | 0.154 | -0.027 | -0.066 |
| **ANXA6** | 0.041 | **0.290** | -0.111 | 0.209 | 0.195 | 0.014 |
| **BGN** | 0.127 | 0.184 | 0.137 | 0.060 | 0.148 | 0.122 |
| **CILP** | 0.010 | 0.132 | -0.012 | -0.154 | 0.118 | -0.020 |
| **CLU** | 0.154 | 0.132 | 0.053 | -0.066 | 0.221 | 0.246 |
| **COL14A1** | 0.166 | 0.012 | 0.058 | -0.178 | 0.033 | 0.188 |
| **COL18A1** | **0.292** | -0.019 | 0.328 | -0.100 | 0.162 | 0.154 |
| **COL6A1** | 0.161 | 0.102 | 0.141 | -0.109 | 0.239 | 0.110 |
| **COL6A2** | 0.215 | 0.035 | 0.214 | -0.045 | 0.213 | 0.122 |
| **COL6A5** | 0.223 | -0.139 | -0.187 | 0.034 | 0.023 | **0.268** |
| **CXCL12** | 0.098 | 0.099 | -0.196 | 0.133 | 0.113 | 0.195 |
| **ELN** | **0.256** | 0.028 | 0.115 | -0.076 | -0.187 | -0.215 |
| **EMILIN1** | **0.345** | -0.011 | 0.189 | 0.006 | **0.310** | **0.298** |
| **FBLN1** | 0.133 | 0.184 | 0.030 | 0.008 | 0.111 | 0.022 |
| **FBLN5** | 0.203 | 0.246 | -0.076 | 0.137 | -0.105 | **-0.257** |
| **FBN1** | 0.121 | 0.088 | -0.035 | -0.143 | 0.087 | 0.107 |
| **GPC6** | -0.054 | **0.297** | -0.222 | 0.000 | 0.006 | 0.043 |
| **HNRNPM** | **0.494** | **-0.455** | 0.149 | 0.138 | 0.101 | **0.427** |
| **LAMB1** | 0.021 | 0.120 | -0.099 | -0.057 | -0.228 | -0.155 |
| **LTBP1** | 0.066 | 0.098 | -0.031 | -0.045 | -0.047 | 0.076 |
| **LUM** | 0.115 | 0.048 | -0.194 | 0.018 | 0.114 | **0.339** |
| **MFAP4** | **0.257** | 0.084 | -0.100 | 0.243 | -0.051 | -0.079 |
| **NID2** | -0.051 | 0.129 | -0.075 | -0.089 | 0.270 | 0.074 |
| **PCOLCE** | 0.134 | 0.005 | 0.082 | -0.133 | **0.398** | **0.394** |
| **PODN** | **0.280** | 0.173 | 0.071 | -0.078 | 0.072 | 0.012 |
| **PRG4** | -0.023 | 0.073 | -0.074 | 0.069 | 0.066 | 0.090 |
| **RARRES2** | 0.238 | 0.176 | -0.029 | 0.209 | -0.002 | -0.005 |
| **SEMA7A** | **0.485** | **-0.298** | 0.076 | 0.213 | 0.220 | **0.501** |
| **SERPINE2** | **0.250** | -0.040 | 0.031 | -0.165 | **0.352** | **0.539** |
| **SERPINF1** | 0.182 | 0.126 | 0.080 | -0.301 | 0.260 | **0.339** |
| **SERPING1** | 0.143 | 0.274 | -0.041 | 0.076 | 0.024 | 0.010 |
| **SMOC2** | **0.262** | 0.165 | 0.124 | -0.031 | 0.001 | 0.088 |
| **SRPX2** | **0.370** | 0.035 | 0.039 | -0.150 | -0.172 | 0.047 |
| **TGFB3** | **0.355** | **-0.355** | 0.010 | 0.103 | 0.221 | **0.471** |
| **TGM2** | **0.499** | -0.115 | 0.068 | 0.117 | 0.206 | **0.297** |
| **THBS1** | **0.809** | **-0.530** | -0.042 | -0.126 | -0.194 | **0.554** |
| **WNT2** | 0.127 | -0.098 | -0.241 | 0.172 | 0.174 | 0.086 |
| **ADAMTSL4** | **-0.281** | -0.013 | 0.087 | -0.057 | 0.112 | **-0.314** |
| **ANGPT4** | -0.157 | -0.054 | 0.142 | 0.133 | -0.099 | **-0.407** |
| **ANOS1** | **-0.321** | 0.043 | -0.088 | **0.546** | 0.071 | **-0.486** |
| **ANXA2** | -0.234 | -0.226 | -0.051 | 0.236 | **0.363** | 0.020 |
| **BCAM** | -0.228 | 0.189 | 0.175 | **0.332** | **0.334** | **-0.285** |
| **CDON** | **-0.306** | 0.104 | 0.090 | -0.010 | -0.141 | -0.201 |
| **COL8A2** | **-0.345** | 0.208 | 0.277 | -0.020 | 0.272 | -0.234 |
| **FREM2** | **-0.287** | -0.202 | 0.021 | -0.010 | -0.003 | **-0.345** |
| **GPC5** | **-0.306** | -0.105 | -0.025 | -0.059 | -0.055 | -0.160 |
| **HAS3** | -0.113 | -0.072 | 0.171 | 0.088 | 0.202 | -0.028 |
| **S100A10** | -0.102 | **-0.309** | -0.124 | **0.369** | 0.122 | 0.060 |
| **SERPINB6** | **-0.392** | 0.217 | -0.030 | -0.073 | -0.239 | **-0.369** |
| **VWA2** | **-0.468** | 0.225 | 0.080 | -0.026 | 0.022 | **-0.450** |

**Table E7: Results of correlations between senescence genes and (SEO-)COPD-associated ECM proteins**

| **R values** | **CDKN1A** | **CDKN1B** | **CDKN2A** | **CDKN2B** | **TP53** | **LMNB1** |
| --- | --- | --- | --- | --- | --- | --- |
| **ANXA6** | 0.073 | -0.129 | 0.067 | -0.104 | -0.164 | 0.050 |
| **CLU** | **0.399** | -0.164 | 0.031 | 0.237 | -0.087 | 0.137 |
| **FBLN5** | **0.412** | -0.108 | 0.074 | 0.115 | -0.097 | 0.000 |
| **MFAP4** | **0.290** | -0.044 | 0.174 | 0.042 | -0.171 | -0.021 |
| **MMP8** | **0.291** | -0.107 | -0.142 | -0.254 | -0.138 | 0.180 |
| **TGM2** | 0.234 | 0.045 | 0.156 | 0.152 | 0.001 | 0.041 |
| **THBS1** | **0.411** | -0.082 | -0.185 | 0.037 | -0.041 | 0.158 |
| **TINAGL1** | 0.084 | 0.059 | 0.136 | 0.243 | 0.080 | -0.017 |
| **CTSH** | -0.193 | 0.007 | 0.165 | 0.001 | -0.113 | -0.236 |
| **LGALS3BP** | -0.209 | 0.047 | 0.256 | -0.064 | 0.121 | -0.004 |
| **PSAP** | -0.011 | 0.060 | 0.159 | -0.081 | 0.090 | 0.014 |

**Table E8: Results of correlations between GSVA scores of senescence signatures and (SEO-)COPD-associated ECM genes**

| **Rho values** | **SenMayo** | **Casella** | **Hernandez** | **Fridman** |
| --- | --- | --- | --- | --- |
| **A2M** | **0.312** | **0.360** | -0.028 | **0.312** |
| **ACAN** | 0.071 | 0.007 | -0.025 | 0.098 |
| **ADAM19** | **0.471** | -0.198 | -0.019 | 0.121 |
| **ADAMTS1** | **0.572** | 0.058 | 0.052 | **0.284** |
| **ADAMTS15** | 0.131 | 0.156 | 0.094 | 0.167 |
| **ADAMTS4** | **0.635** | 0.023 | 0.189 | **0.311** |
| **ADAMTS8** | **0.458** | 0.155 | **0.440** | **0.379** |
| **ANGPT1** | 0.057 | 0.037 | **-0.300** | -0.028 |
| **ANXA6** | **0.244** | **0.272** | -0.074 | **0.333** |
| **BGN** | **0.289** | 0.219 | 0.046 | **0.395** |
| **CILP** | -0.006 | 0.024 | -0.208 | -0.023 |
| **CLU** | **0.252** | **0.308** | 0.126 | **0.295** |
| **COL14A1** | 0.038 | -0.019 | -0.254 | -0.045 |
| **COL18A1** | **0.367** | **0.277** | 0.264 | **0.431** |
| **COL6A1** | **0.262** | **0.296** | 0.057 | **0.357** |
| **COL6A2** | **0.336** | **0.358** | 0.139 | **0.422** |
| **COL6A5** | 0.218 | -0.081 | -0.060 | 0.001 |
| **CXCL12** | 0.194 | 0.037 | -0.163 | 0.129 |
| **ELN** | 0.190 | 0.141 | 0.095 | **0.262** |
| **EMILIN1** | **0.537** | **0.359** | 0.256 | **0.565** |
| **FBLN1** | **0.242** | **0.426** | 0.042 | **0.385** |
| **FBLN5** | 0.217 | 0.234 | 0.083 | **0.352** |
| **FBN1** | 0.172 | 0.143 | -0.094 | 0.156 |
| **GPC6** | -0.026 | -0.078 | **-0.382** | -0.061 |
| **HNRNPM** | **0.527** | **0.257** | **0.494** | **0.569** |
| **LAMB1** | 0.030 | 0.132 | -0.081 | 0.140 |
| **LTBP1** | 0.063 | 0.167 | -0.149 | 0.141 |
| **LUM** | **0.236** | 0.114 | -0.259 | 0.095 |
| **MFAP4** | **0.352** | **0.311** | 0.167 | **0.414** |
| **NID2** | 0.101 | 0.013 | -0.051 | 0.036 |
| **PCOLCE** | **0.286** | **0.277** | 0.031 | **0.354** |
| **PODN** | **0.309** | **0.319** | 0.103 | **0.391** |
| **PRG4** | 0.063 | 0.150 | -0.164 | 0.100 |
| **RARRES2** | **0.317** | 0.158 | 0.045 | **0.305** |
| **SEMA7A** | **0.543** | 0.084 | **0.347** | **0.364** |
| **SERPINE2** | **0.360** | 0.151 | -0.063 | 0.231 |
| **SERPINF1** | **0.283** | **0.278** | -0.079 | **0.237** |
| **SERPING1** | **0.248** | **0.353** | -0.073 | **0.324** |
| **SMOC2** | **0.256** | 0.164 | -0.018 | **0.250** |
| **SRPX2** | **0.252** | 0.005 | 0.010 | 0.141 |
| **TGFB3** | **0.472** | 0.091 | 0.159 | **0.289** |
| **TGM2** | **0.657** | **0.281** | 0.272 | **0.520** |
| **THBS1** | **0.722** | 0.025 | 0.207 | **0.365** |
| **WNT2** | **0.316** | **0.292** | 0.108 | **0.345** |
| **ADAMTSL4** | **-0.263** | 0.185 | 0.211 | -0.059 |
| **ANGPT4** | -0.111 | 0.199 | **0.341** | 0.011 |
| **ANOS1** | -0.222 | 0.083 | 0.177 | 0.003 |
| **ANXA2** | -0.057 | **0.332** | 0.264 | 0.173 |
| **BCAM** | -0.088 | **0.389** | **0.401** | **0.317** |
| **CDON** | **-0.417** | -0.072 | -0.067 | **-0.292** |
| **COL8A2** | -0.204 | **0.353** | 0.118 | 0.025 |
| **FREM2** | **-0.418** | -0.153 | 0.042 | **-0.354** |
| **GPC5** | **-0.410** | -0.170 | 0.010 | **-0.367** |
| **HAS3** | -0.057 | -0.036 | 0.130 | -0.134 |
| **S100A10** | 0.029 | 0.094 | 0.142 | 0.108 |
| **SERPINB6** | **-0.560** | **-0.388** | -0.185 | **-0.511** |
| **VWA2** | **-0.458** | -0.054 | 0.068 | **-0.243** |

**Table E9: Results of correlations between GSVA scores of senescence signatures and (SEO-)COPD-associated ECM proteins**

| **Rho values** | **SenMayo** | **Casella** | **Hernandez** | **Fridman** |
| --- | --- | --- | --- | --- |
| ANXA6 | 0.017 | -0.066 | -0.052 | -0.027 |
| CLU | **0.268** | -0.050 | 0.162 | 0.222 |
| FBLN5 | **0.325** | 0.070 | 0.166 | **0.287** |
| MFAP4 | 0.210 | 0.020 | 0.094 | 0.148 |
| MMP8 | **0.376** | -0.100 | -0.014 | 0.105 |
| TGM2 | 0.207 | 0.011 | 0.082 | 0.120 |
| THBS1 | **0.419** | 0.037 | 0.203 | **0.332** |
| TINAGL1 | 0.060 | 0.017 | 0.050 | 0.111 |
| CTSH | -0.235 | -0.061 | 0.034 | -0.176 |
| LGALS3BP | -0.067 | 0.000 | 0.143 | -0.050 |
| PSAP | 0.046 | 0.020 | 0.198 | 0.047 |

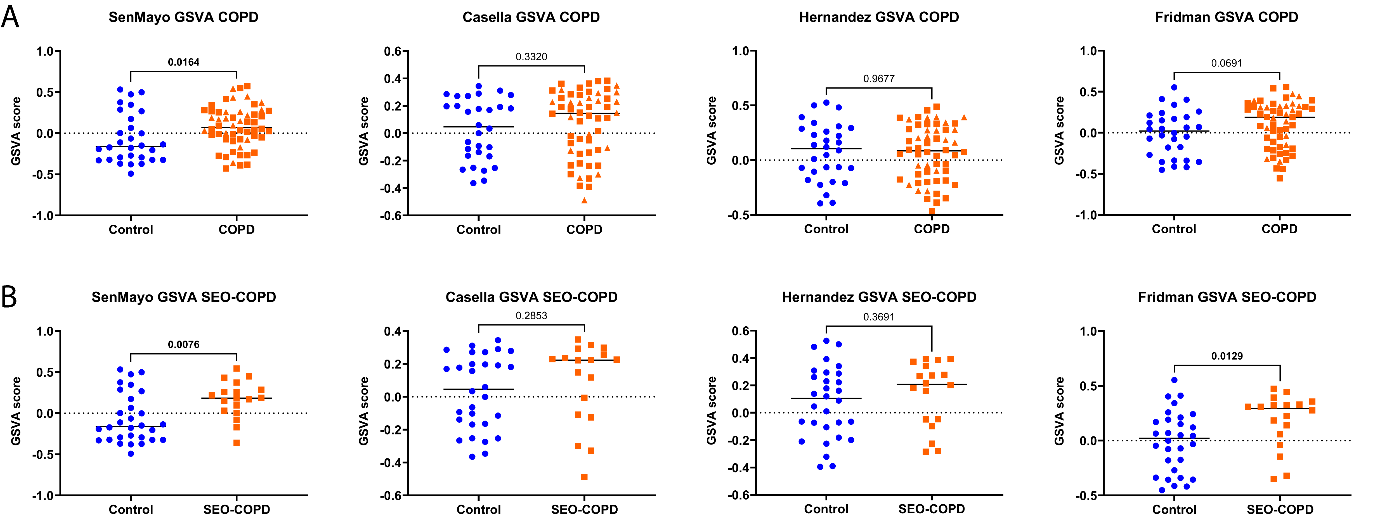

***Figure E1: Higher GSVA score of senescence signatures in (SEO-)COPD lung tissue.*** *GSVA score was calculated using the gene expression in lung tissue from (SEO-)COPD patients and controls for the genes in the senescence signatures; SenMayo, Casella, Hernandez-Segura, and Fridman. GSVA scores were compared between COPD and control (A) and SEO-COPD and control (B) using Mann Whitney U tests. P-values are indicated and bold when significant (P<0.05).*

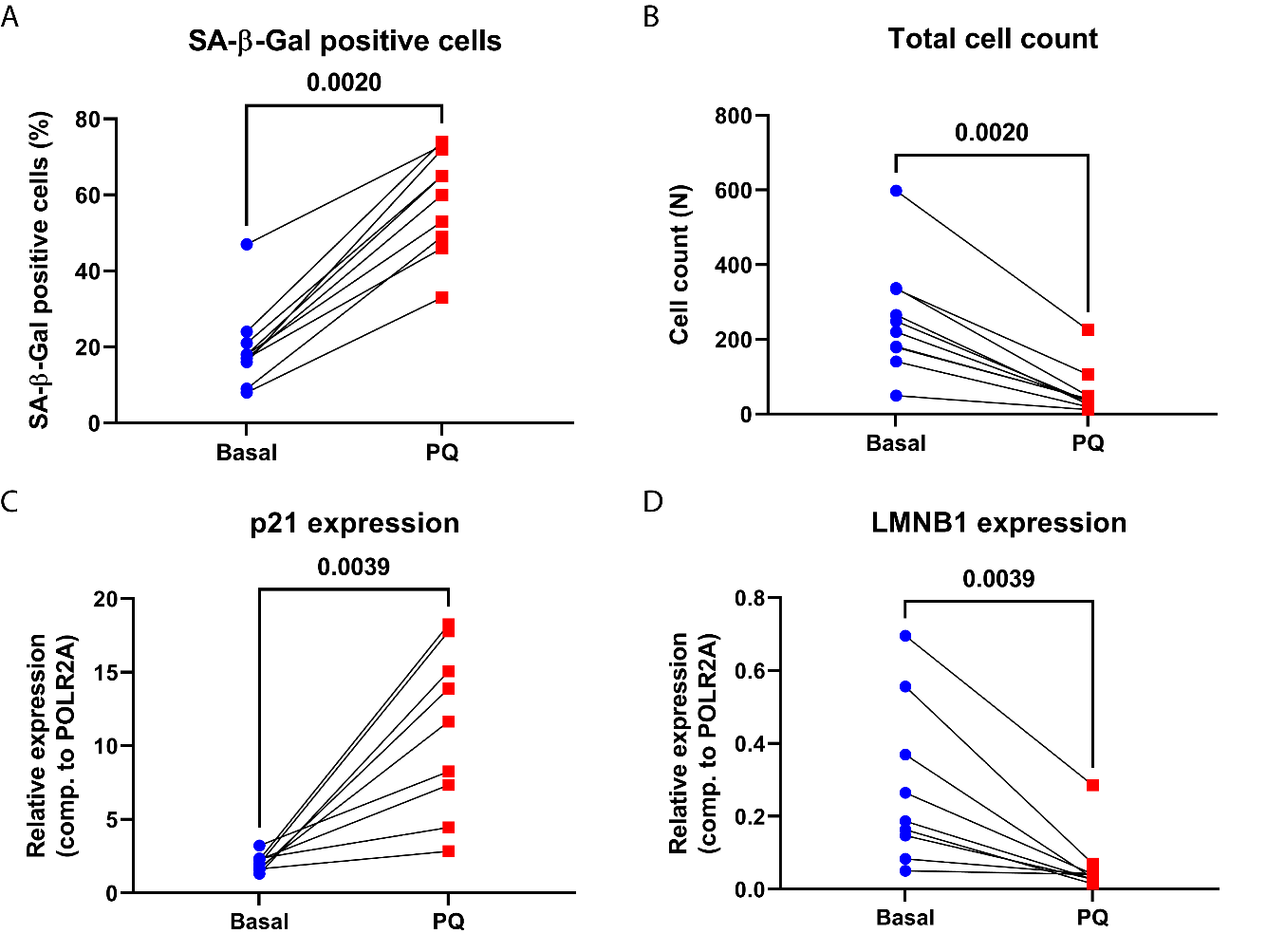

***Figure E2: Cellular senescence induction in primary lung fibroblasts.*** *Senescence was induced in primary lung fibroblasts with 250uM PQ and after 5 days cells were reseeded to reach same confluency and SA-β-gal staining was performed and gene expression was measured using RT-qPCR after 3 days of reseeding. The percentage of SA-β-gal positive cells (A) and total cell counts (B) are depicted in the dot plots on top. Gene expression of CDKN1A (C) and LMNB1 (D) of untreated (Basal, blue) and PQ-induced senescent (PQ, red) fibroblasts are shown in the dot plots below. The lines represent matching donors. Relative expression was calculated using POLR2A as housekeeping gene and depicted. Statistical significance was tested using Wilcoxon signed-rank test, p-values are indicated and bold when < 0.05.*
